# Transcriptome mining of RNA viruses (family *Totiviridae*) in *Eimeria necatrix* and *Eimeria stiedai*

**DOI:** 10.1101/2023.05.20.541574

**Authors:** Max L. Nibert, Yue Xie, Jie Xiao, Yang Gao, Dandan Liu, Guangyou Yang, Jianping Tao

**Author notes:** Corresponding authors: mnibert.hms.harvard.edu.

## Abstract

Coccidian protozoa from the genus *Eimeria* are widespread parasites of vertebrates, causing serious disease (coccidiosis) and economic loss most notably in poultry. Several species of *Eimeria* are themselves infected with small RNA viruses assigned to the family *Totiviridae*. In this study, the sequences of two such viruses were newly determined, one of which represents the first complete protein-coding sequence of a virus from *E. necatrix*, an important pathogen of chickens, and the other of which is from *E. stiedai*, an important pathogen of rabbits. Sequence features of the newly identified viruses, compared with those of ones reported previously, provide several insights. Phylogenetic analyses suggest that these eimerian viruses constitute a well-demarcated clade, probably deserving of recognition as a distinct genus.

## INTRODUCTION

Members of the genus *Eimeria* are protozoan parasites (Apicomplexa; Conoidasida; Coccidia) that cause enteric or hepatic disease in a wide variety of vertebrates (Chapman et al., 2013; Lee et al., 2022; López-Osorio et al., 2020). Importantly, they are associated with serious illness (coccidiosis) in several economically important food animals such as cattle, sheep, goats, rabbits, chickens, and turkeys. *Eimeria necatrix* causes avian coccidiosis and is one of the most virulent coccidian pathogens of chickens (Gao et al., 2021). To date, at least nine additional species have been implicated in chicken coccidiosis, including *E. acervulina, E. brunetti, E. maxima, E. mitis, E. praecox*, and *E. tenella* (Blake et al., 2021; Gerhold, 2023). *Eimeria stiedai*, on the other hand, causes rabbit coccidiosis and is the most virulent coccidian pathogen of rabbits (Xie et al., 2021). To date, fourteen additional species have been implicated in rabbit coccidiosis (Xie et al., 2021), including *E. flavescens, E. intestinalis, E. irresidua, E. magna, E. media*, and *E. perforans*. Transmission of eimerian coccidia occurs largely by the fecal–oral route, and typical symptoms include diarrhea, dehydration, weight loss, lethargy, and growth retardation, sometimes followed by death.

Eimerian coccidia have complex life cycles, with an exogenous phase in the environment and an endogenous phase in the vertebrate host (Gao et al., 2021; López-Osorio et al., 2020; Xie et al., 2021). Oocysts excreted into the environment during host infection undergo meiosis and cell division to form sporozoites (sporogony). After ingestion by the next host individual, the sporocyst wall is disrupted by mechanical and biochemical forces along the host gastrointestinal tract, leading to release of the sporozoites in the small intestine and migration to their preferred site for cellular invasion. Upon entering the host cell, each sporozoite transforms to a feeding stage called a trophozoite. The cycle next proceeds through repeated trophozoite, schizont (or meront), and merozoite stages, which is known as schizogony (or merogony). Second- or third-generation merozoites then go on to form the gametocyte stages (gametogony) that mediate fertilization and produce the next generation of oocysts for excretion in feces. Rupture of host cells during schizogony/merogony and gametogony is thought to be a major contributor to disease, though pro-inflammatory host responses may also play a role (Lee et al., 2022). Detailed molecular aspects of eimerian interactions with host cells remain poorly understood.

One interesting aspect of eimerian biology is that several species are naturally infected with small RNA viruses assigned to the family *Totiviridae*, probably in a persistent and largely noncytopathic manner as known for other *Totiviridae* members (Goodman et al., 2011; Wickner et al., 2011). To date, the presence of such viruses in isolates of *E. brunetti, E. necatrix, E. nieschulzi, E. stiedai*, and *E. tenella* seems fairly well established (del Cacho et al., 2001; Ellis and Revets, 1990; Han et al., 2011; Lee and Fernando, 1998, 1999, 1999b, 2000; Lee et al., 1996; Revets et al., 1989; Roditi et al., 1994; Sepp et al., 1991; Wang et al., 2018; Wu et al., 2016; Xin et al., 2016), with complete protein-coding sequences reported for viruses from *E. brunetti, E. stiedai*, and *E. tenella* (e.g., Reference Sequence (RefSeq) accessions NC_026140.1, NC_002701.1, and NC_040530.1; Wu et al., 2016; Xin et al., 2016) and short partial sequences for viruses from *E. necatrix* (Lee et al., 1998) and rat pathogen *E. nieschulzi* (GenBank accession L25869.1; Roditi et al., 1994).

In studies for this report, we sought to obtain complete protein-coding sequences for other eimerian viruses, in particular by mining transcriptome datasets, including RNA-derived Sequence Read Archive (SRA) libraries, that are publicly available at GenBank and the National Center for Biotechnology Information (NCBI; Bethesda, MD, USA). As described in detail below, we succeeded in assembling such sequences by using the SRA libraries from defined laboratory isolates of two *Eimeria* species, one from *E. necatrix* isolate Yangzhou and another from *E. stiedai* isolate SCES/Sichuan.

## MATERIALS AND METHODS

### Sequence assemblies

Steps in the assembly process for the two newly identified eimerian viruses are summarized in Results. For the virus from *E. necatrix* isolate Yangzhou, the following RNA-derived SRA libraries from BioProjects PRJNA730226 and PRJNA730346 (Gao et al., 2021) were used: SRX10900611– SRX10900613 (150-bp Illumina reads from sporozoites), SRX10900614–SRX10900616 (150-bp Illumina reads from unsporulated oocysts), SRX10904653 (PacBio reads from unsporulated oocysts), and SRX10904654 (PacBio reads from sporozoites). Only the Illumina reads were used in generating the final draft, refined, and de novo assembled sequences. For the virus from *E. stiedai* isolate SCES/Sichuan, the following RNA-derived SRA libraries from BioProject PRJNA730346 (Xie et al., 2021) were used: SRX9985032, SRX9985033, and SRX9985037 (150-bp Illumina reads from sporulated oocysts) and SRX9985038–SRX9985040 (150-bp Illumina reads from unsporulated oocysts).

Tools used in generating the sequence assemblies were largely those as implemented at Galaxy Australia (https://usegalaxy.org.au/). For CAP3 (v. 2.0.0), an overlap length cutoff of 16 and an overlap percent identity cutoff of 66 were used. For smaller numbers of reads, CAP3 as implemented at https://doua.prabi.fr/software/cap3 was used. For Bowtie2 (v. 2.5.0+galaxy0), default settings were used with a reference from the history and paired-end reads as a dataset collection from the SRA libraries. For iVar consensus (v. 1.4.0+galaxy0), default settings were used on the output from Bowtie2 except with a minimum frequency of 50% and a minimum depth of 1. When a minimum frequency of 80% was used as a test, the assemblies for EnecRV1-Yangzhou and EstiRV1-SCES were found to include only one or four ambiguous residues, respectively, attesting to their robustness. For rnaviralSPAdes (v. 3.15.4+galaxy2), default settings were used with paired-end reads as a list of dataset pairs from the SRA libraries. Several auxiliary programs used in the overall process included SAMtools Idxstats for obtaining read numbers and deepTools plotCoverage for analyzing positional coverage depths. Different options from the BLAST suite, as indicated in this report, were generally used as implemented at https://blast.ncbi.nlm.nih.gov/, except for searches of rnaviralSPAdes libraries, which was done at Galaxy Australia using NCBI BLAST+ options.

### Phylogenetic analyses

Curation of the RefSeq accessions obtained using the NCBI Virus resource for Fig. 2 included elimination of one apparently misclassified sequence (NC_025367.1), elimination of one sequence (NC_005965.1) that shared >90% identity with another sequence (NC_003876.1), and correction of what appears to be a truncated RdRp sequence from accession NC_005883.1. Curation of the RefSeq and regular GenBank accessions used for Fig. 3 is described in Results. Alignments were performed with MAFFT-L-INS-I (v. 7.511) as implemented at https://mafft.cbrc.jp/alignment/server/ (Katoh et al., 2019; default settings except offset value 0.123). Alignments were then curated with TrimAl (v. 1.4.1) (Capella-Gutiérrez et al., 2009) as implemented at https://ngphylogeny.fr/ (Lemoine et al., 2019; automatic option). Maximum-likelihood phylogenetic analyses were performed with IQ-TREE (v. 1.6.11) as implemented at https://www.hiv.lanl.gov/content/sequence/IQTREE/iqtree.html(Trifinopoulos et al., 2016; ‘find best and apply’ option for Substitution Model, 1000 ultrafast bootstraps for Branch Support, default settings for Tree Search and rooting options). Trees were prepared for publication using FigTree (v. 1.4.2). Elements specific to each tree, such as the best-fit substitution model determined and applied by IQ-TREE, are indicated in the respective figure legends.

**Fig. 1.**
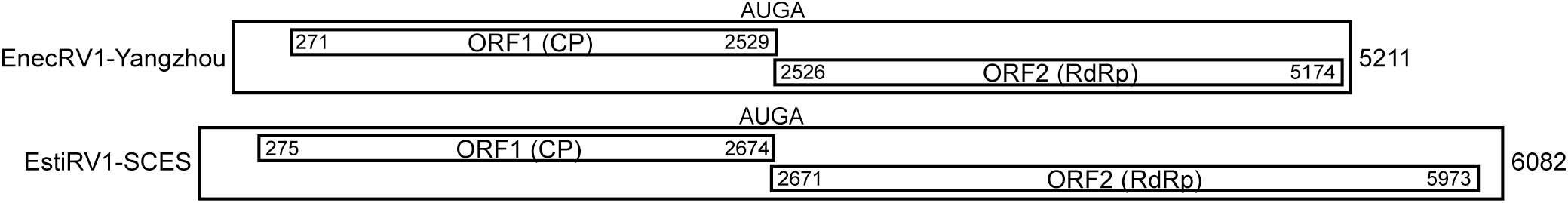
Genome diagrams. The newly reported nt sequences of EnecRV1-Yangzhou and EstiRV1-SCES are diagrammed to scale, with positions numbered at the end of each sequence and the beginning and end of each ORF. Position of the AUGA stop/restart motif is indicated at the end of ORF1 and beginning of ORF2 in each sequence.

**Fig. 2.**
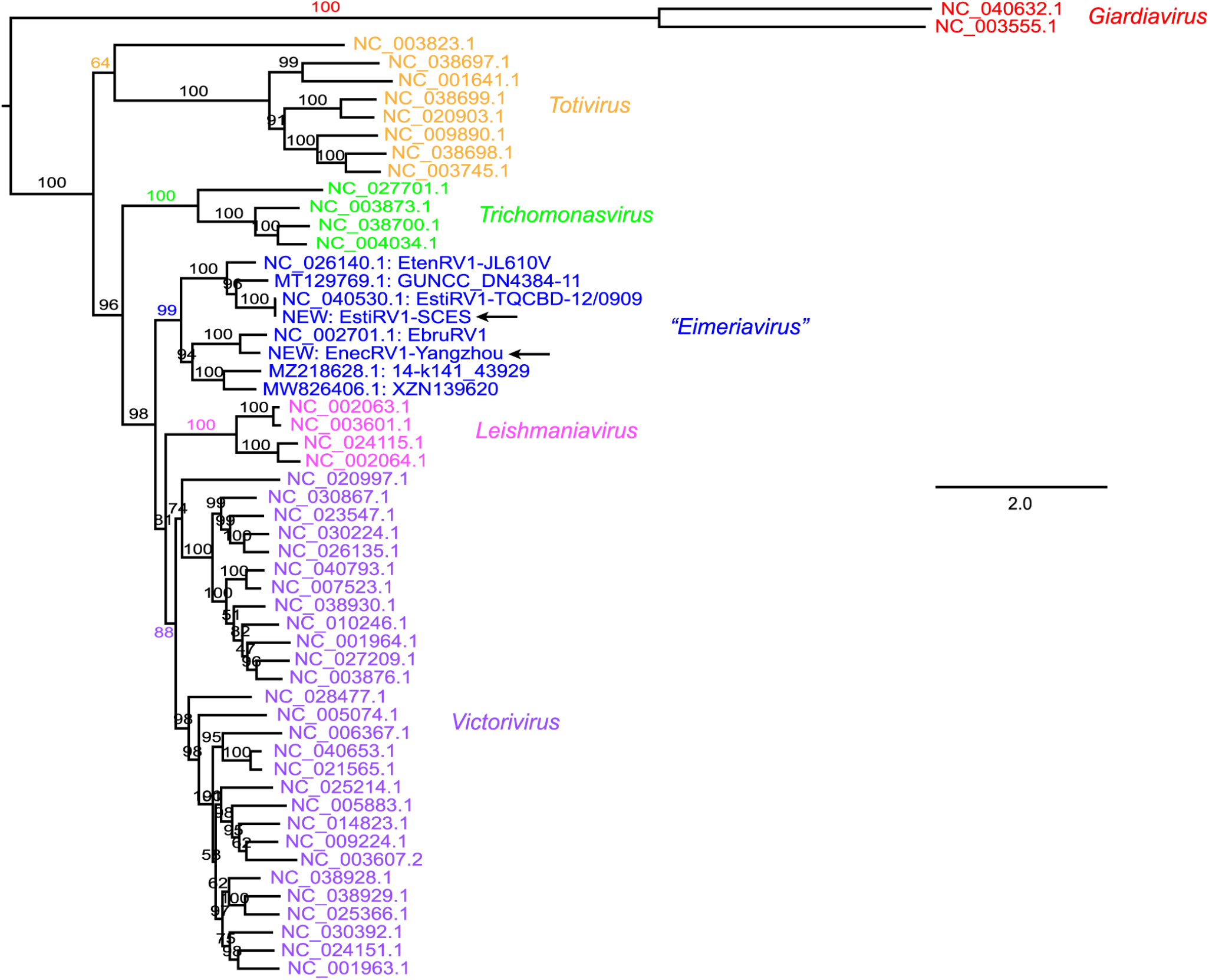
Phylogenetic tree 1. As explained in the main text, these analyses included 46 RdRp sequences translated from RefSeq accessions (see branch labels) assigned to genus *Giardiavirus* (red), *Totivirus* (orange), *Trichomonasvirus* (green), *Leishmaniavirus* (magenta), or *Victorivirus* (purple) at NCBI, along with 8 RdRp sequences from putative eimerian viruses with coding-complete sequences (blue). After trimming with trimAl, the MAFFT-L-INS-i alignment of these sequences spanned 654 aa positions. The best-fit substitution model according to the Bayesian information criterion, as determined by ModelFinder within IQ-TREE, was LG+F+R5 (proportion of invariant sites, 0.0260; model of rate heterogeneity, FreeRate with 5 categories; site proportion and rates, (0.1175, 0.0965) (0.1760, 0.3266) (0.2472, 0.6856) (0.2436, 1.1767) (0.2158, 2.2018)). Branch support values as determined by UFBoot2 within IQ-TREE are shown in %. The tree is rooted on the midpoint, showing the greater divergence of the giardiavirus clade as observed previously (Janssen et al., 2015). Scale bar indicates average number of substitutions per trimmed alignment position. Arrows highlight the new sequences in this report.

**Fig. 3.**
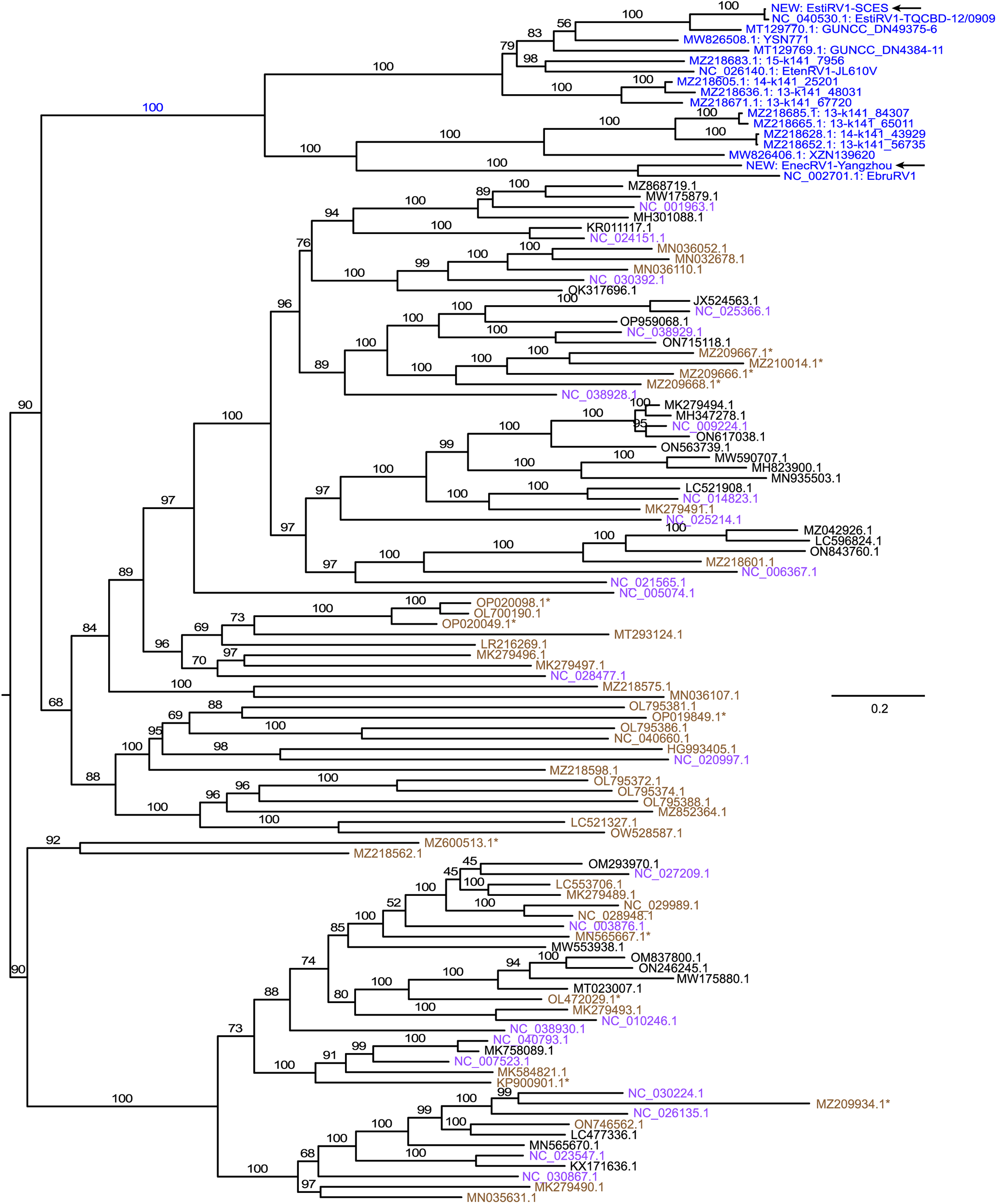
Phylogenetic tree 2. As explained in the main text, these analyses included 113 unique RdRp sequences obtained through BLASTp searches using the RdRp sequences of the 8 putative eimerian viruses with coding-complete nt sequences. The RdRp sequences of the two newly reported viruses in this report were added for a total of 115. RefSeq and GenBank accession numbers have been converted to those from which the RdRp sequences were translated. Color coding: blue, sequences attributed to eimerian viruses; purple, RefSeq accessions assigned to genus *Victorivirus* at NCBI and appearing in Fig. 2; black, GenBank accessions assigned to genus *Victorivirus* at NCBI and not appearing in Fig. 2; and brown, RefSeq or GenBank accessions not assigned to genus *Victorivirus* at NCBI, not appearing in Fig. 2, and not mapping to the eimerian virus clade. The 12 accessions in brown and marked with an asterisk appear to be incorrectly assigned to genus *Totivirus* at NCBI; the other accessions in brown are assigned to family *Totiviridae*, but not to a genus. After trimming with trimAl, the MAFFT-L-INS-i alignment of these sequences spanned 656 aa positions. The best-fit substitution model according to the Bayesian information criterion, as determined by ModelFinder within IQ-TREE, was LG+F+R8 (proportion of invariant sites, 0.0732; model of rate heterogeneity, FreeRate with 8 categories; site proportion and rates, (0.1122, 0.0220) (0.0576, 0.0958) (0.1091, 0.2526) (0.1373, 0.4897) (0.1998, 0.7709) (0.2016, 1.3304) (0.1093, 2.1821) (0.0730, 3.2381)). Branch support values as determined by UFBoot2 within IQ-TREE are shown in %. The tree is rooted on the midpoint. Scale bar indicates average number of substitutions per trimmed alignment position. Arrows highlight the new sequences in this report.

## RESULTS

### RNA virus in chicken pathogen *Eimeria necatrix*

We applied Discontiguous MegaBLAST to search the RNA-derived SRA libraries from unsporulated oocysts and sporozoites of *E. necatrix* isolate Yangzhou (BioProjects PRJNA730226 and PRJNA730346; Gao et al., 2021), using previously reported virus sequences as queries. Query accession NC_002701.1 for Eimeria brunetti RNA virus 1 (EbruRV1) provided a large number of hits, including ones from all six SRA libraries containing Illumina reads and both SRA libraries containing PacBio reads. Illumina-read hits were then used with CAP3 to assemble an apparently coding-complete final draft sequence, 5,211 nt in length, of what appeared to be the genome of a newly identified virus. In particular, this draft sequence was notable for containing two long open reading frames (ORFs), respectively encoding the capsid protein (CP) and the RNA-dependent RNA polymerase (RdRp) of a probable *Totiviridae* member based on BLASTp searches with both products. The draft sequence was next subjected to refinement by using it as reference with Bowtie2 to remap reads from the six SRA libraries, followed by reassembly with iVar Consensus to obtain a refined sequence from the mapped reads. After resolving minor differences at the 3′-terminus, the refined sequence was found to be identical to the draft, except for a single A-to-G transition, and was calculated to have an RPKM of 1.1 and an average coverage depth of 71. Lastly, fully *de novo* assembly with rnaviralSPAdes was applied to the six SRA libraries containing short reads to rule out any bias that might have been present during the preceding “map-reads-to-reference” approaches. A MegaBLAST search of this *de novo* assembled transcriptome of *E. necatrix* isolate Yangzhou, using the refined sequence as query, identified a single hit, which was found to be identical to the refined sequence. This corroborated sequence has been deposited in GenBank as accession OQ818109 under the name Eimeria necatrix RNA virus 1 strain Yangzhou (EnecRV1-Yangzhou).

### RNA virus in rabbit pathogen *Eimeria stiedai*

Essentially the same series of steps was used on the RNA-derived SRA libraries from *E. stiedai* isolate SCES/Sichuan (BioProject PRJNA698271; Xie et al., 2021). In this case, a large number of hits from the initial search was obtained using query accession NC_040530.1 for Eimeria stiedai RNA virus 1 isolate TQCBD-12/0909 (EstiRV1-TQCBD-12/0909) (Xin et al., 2016). Interestingly, hits were obtained with all six SRA libraries containing Illumina reads from unsporulated and sporulated oocysts, but not with those from merozoites or gametophytes. The Illumina-read hits from unsporulated and sporulated oocysts were then used to assemble an apparently coding-complete final draft sequence, 6,082 nt in length, of what appeared to be the genome of another newly identified virus. This draft sequence was again notable for containing two long ORFs, respectively encoding the CP and the RdRp of an apparent *Totiviridae* member based on BLASTp searches with both products. The sequence then subjected to refinement by remapping and reassembly was found to be identical to the reference draft, except for a single G-to-A transition, and was calculated to have an RPKM of 1.7 and a median coverage depth of 84. Lastly, the virus sequence extracted from the *de novo* assembled transcriptome of *E. stiedai* isolate SCES/Sichuan was found to be identical to the refined sequence. This corroborated sequence has been deposited in GenBank as accession OQ821982 under the name Eimeria stiedai RNA virus 1 strain SCES (EstiRV1-SCES).

### Basic sequence features of the newly found viruses

The 5,211-nt sequence of EnecRV1-Yangzhou includes a 270-nt nontranslated region (NTR) at its 5′ terminus and a 37-nt NTR at its 3′ terminus. In between, the CP-encoding ORF, ORF1, spans residues 271–2,529 from its first in-frame AUG codon to its UGA stop codon, and the RdRp-encoding ORF, ORF2, spans residues 2,526–5,174 from its first in-frame AUG codon to its UAA stop codon. The small overlap between these two ORFs comprises the ORF2 start codon and the ORF1 stop codon in the sequence AUGA. This sequence suggests use of a stop/restart mechanism for translation of the downstream ORF, as described for members of the genus *Victorivirus* in family *Totiviridae* (Li et al., 2011), as well as for members of other virus families (Firth and Brierley, 2012). The nature of this motif in other eimerian viruses is addressed in Discussion. The top-scoring hit to the EnecRV1-Yangzhou sequence in a BLASTn search was accession NC_002701.1 for EbruRV1 (E-value, 3e-129; query cover, 70%; identity, 66%), i.e., the same accession used as reference query in the discovery of this virus. A global alignment of these two nt sequences using EMBOSS Needle yielded an identity score of only 59%, suggesting that EnecRV1-Yangzhou may be considered to represent a new virus species. EbruRV1 is currently identified at NCBI as an unclassified member of the family *Totiviridae*. For comparison, the length of the EbruRV1 genome is reported as 5,358 nt, with a 357-nt 5′ NTR and a 37-nt 3′ NTR flanking its two ORFs, which themselves overlap in the stop/restart motif AUGA.

The 6,082-nt sequence of EstiRV1-SCES includes a 274-nt NTR at its 5′ terminus and a 110-nt NTR at its 3′ terminus. In between, the CP-encoding ORF, ORF1, spans residues 275–2,674 from its first in-frame AUG codon to its UGA stop codon, and the RdRp-encoding ORF, ORF2, spans residues 2,671–5,973 from its first in-frame AUG codon to its UAG stop codon. The small overlap between these two ORFs again comprises the ORF2 start codon and the ORF1 stop codon overlapping in the sequence AUGA, a putative stop/restart motif for translation of the downstream ORF. The top-scoring hit to the EstiRV1-SCES sequence in a BLASTN search was accession NC_040530.1 for EstiRV1-TQCBD-12/0909 (E-value, 0.0; query cover, 100%; identity, 99%), i.e., the same accession used as reference query in the discovery of this virus. A global alignment of these two nt sequences using EMBOSS Needle yielded an identity score of 96%, suggesting that EstiRV1-SCES does not represent a new virus species, but rather another strain of the same species as EstiRV1-TQCBD-12/0909. As noted above for EbruRV1, EstiRV1 too is currently identified at NCBI as an unclassified member of the family *Totiviridae*. For comparison, the length of the EstiRV1-TQCBD-12/0909 genome is reported as 6,219 nt, with a 391-nt 5′ NTR and a 129-nt 3′ NTR flanking its two ORFs, which themselves overlap in the stop/restart motif AUGA.

Based on BLAST searches, ORF1 and ORF2 in each new sequence are concluded to encode the viral CP and the viral RdRp, respectively. Deduced length of the CP is 752 aa in EnecRV1-Yangzhou and 799 aa in EstiRV1-SCES. These compare with CP lengths of 770 aa in EbruRV1 and 799 aa in EstiRV1-TQCBD-12/0909. Deduced length of the RdRp is 882 aa in EnecRV1-Yangzhou and 1,100 aa in EstiRV1-SCES. These compare with RdRp lengths of 884 aa in EbruRV1 and 1,100 aa in EstiRV1-TQCBD-12/0909. The identical lengths of the CP and RdRp in EstiRV1-SCES and EstiRV1-TQCBD-12/0909 reflect their high degree of relatedness. Identity scores in pairwise comparisons using EMBOSS Needle are 99.1% for EstiRV1-SCES and EstiRV1-TQCBD-12/0909 versus only 59.9% for EnecRV1-Yangzhou and EbruRV1 when comparing the concatenated CP and RdRp sequences of each pair.

### Review of apparent eimerian virus sequences in GenBank

In preparation for phylogenetic analyses, BLASTx searches, using the two new virus nt sequences as queries, pointed to six GenBank accessions that represent what appear to be coding-complete sequences of eimerian viruses. Three of the six accessions are annotated as viruses of particular *Eimeria* species and indeed are also represented by RefSeq accessions (EbruRV1, AF356189.1 and NC_002701.1; EstiRV1-TQCBD-12/0909, KU597305.1 and NC_040530.1; and Eimeria tenella RNA virus 1 strain JL610V (EtenRV1-JL610V), KJ363185.1 and NC_026140.1). These include the two reference queries used in discovering the two new viruses. One other of the six accessions, though annotated as being associated with *E. stiedai*, was obtained from pooled cecal contents of wild rabbits (Eimeria stiedai RNA virus 1 strain GUNCC_DN54384-11, MT129769.1) without evidence implicating *E. stiedai* in particular. The remaining two of the six accessions are designated simply as Totiviridae sp. but were obtained from feces of pigeons (isolate 14-k141_43929, MZ218628.1) or other wild birds (isolate XZN139620, MW826406.1). The BLASTx searches also pointed to ten GenBank accessions that represent what appear to be partial sequences of other eimerian viruses. These partial sequences were obtained from samples of pooled organ tissues of wild bats (MN851287.1) (Kohl et al., 2021), pooled cecal contents of wild rabbits (MT129770.1) (Mahar et al., 2020), or feces of pigeons (MZ218605.1, MZ218636.1, MZ218652.1, MZ218665.1, MZ218671.1, MZ218683.1, and MZ218685.1; Chen et al., 2022) or other wild birds (MW826508.1; Zhu et al., 2022), and indeed the first two of the ten accessions (MN851287.1 and MT129770.1) are annotated as being associated with *Eimeria* species (the others again as simply Totiviridae sp.). Since members of genus *Eimeria* are widespread enteric pathogens of vertebrates, commonly transmitted by the fecal-oral route, evidence for eimerian viruses in wild-collected samples such as rabbit cecal contents or bird feces is not unexpected.

In an effort to clarify which *Eimeria* species may have been infected with the apparent eimerian viruses that lack clearly identified host species, we examined mitochondrial genetic markers that might provide clues in that regard, in particular transcripts encoding proteins cytochrome b (cytb) and cytochrome c oxidases 1 and 3 (cox1 and cox3) (Zhou et al., 2023). We first identified reads representing these transcripts from the SRA library (SRX7914142) from which the Eimeria stiedai RNA virus 1 GUNCC strains from rabbit cecal contents were derived (MT129769.1 and MT129770.1). These reads were assembled into contigs spanning 1235, 1432, and 777 bp, respectively, which were in turn used to search against the *Eimeria* mitochondrial sequences available in GenBank. The results suggest these viruses may have come from one or more other rabbit-infecting species—in particular, *E. magna* (KF419217.1; Tian et al., 2015; nt identity score for *cytb+cox1+cox3*, 99.5%; aa identity score, 100%) and/or *E. media* (KP025691.1; nt identity score for *cytb+cox1+cox3*, 99.0%)—instead of *E. stiedai* (OQ427352.1; nt identity scores for *cytb+cox1+cox3*, 97.6%). The same approach was applied to the SRA libraries (SRX10416136, SRX10416147, and SRX10416158) from which the Totiviridae sp. strains from pigeon feces were derived (MZ218628.1, MZ218605.1, MZ218636.1, MZ218652.1, MZ218665.1, MZ218671.1, MZ218683.1, and MZ218685.1). Several *Eimeria* species are known to infect pigeons, *e*.*g*., *E. columbarum* and *E. labbeana*, but none of those are well represented in GenBank to date. Nonetheless, reads from each of the three SRA accessions were assembled into contigs spanning 1076–1240, 1414–1441, and 732–830 bp for cytb-, cox1-, and cox3-encoding transcripts, respectively, and the search results suggest these viruses may have come from one or more *Eimeria* species most closely (though still somewhat distantly) related to turkey-infecting species *E. innocua* (KR108296.1; Ogedengbe et al., 2014; nt identity scores for *cytb+cox1+cox3*, 95.3–95.7%) and *E. dispersa* (KJ608416; Hafeez et al., 2016; nt identity scores for *cytb+cox1+cox3*, 95.0–95.4%). Although these findings are not definitive due to the metagenomic nature of the samples, they point toward several other *Eimeria* species that may be infected with closely related small RNA viruses.

### Phylogenetic analyses

Sequences previously attributed to eimerian viruses have been found to form a discrete phylogenetic cluster, positioned near mycovirus members of the genus *Victorivirus* and have been proposed to be assigned either to genus *Victorivirus* itself, perhaps as a distinct subgenus (Wu et al., 2016), or instead to a new genus with proposed name “Eimeriavirus” (Xin et al., 2016). For phylogenetic analyses in this study, we used the NCBI Virus resource (https://www.ncbi.nlm.nih.gov/labs/virus/) to download all RefSeq genome accessions for viruses assigned to the genera *Victorivirus, Leishmaniavirus, Trichomonasvirus, Totivirus*, and *Giardiavirus* in family *Totiviridae*. After curation, a set of 46 RdRp sequences was translated from these accessions. The RdRp sequences of the eight apparent eimerian viruses with coding-complete sequences, including the two new viruses, were then added to this set before being aligned with MAFFT-L-INS-i, trimmed with TrimAl, and subjected to maximum-likelihood phylogenetic analyses with IQ-TREE. In the resulting trees, the RdRp sequences of the eight apparent eimerian viruses consistently defined a discrete clade with strong branch support (Fig. 2), in line with previously reported findings (Chen et al., 2022; Wu et al., 2016; Xin et al., 2016). Moreover, positionally, the clade comprising the eimerian viruses was sandwiched between a discrete clade comprising the trichomonasviruses and a larger clade comprising both the leishmaniaviruses and the victoriviruses in discrete subclades of their own. The same features were seen when these analyses also included the ten partial sequences of likely other eimerian viruses described above.

To challenge these results by including other closely related viruses in the analyses, we obtained a second set of RdRp sequences by using the eight apparent eimerian viruses with coding-complete sequences as queries in BLASTp searches. The top 100 hits based on E-values from each search were then downloaded, after which they were combined, duplicates were eliminated, and sequences with >90% identity to another sequence in the set were also eliminated, yielding a set of 113 unique RdRp sequences, 115 when those of the two new viruses were added. Seventeen of the 115 sequences were identified as belonging to the 18 apparent eimerian viruses discussed in the first paragraph of this section; only the partial RdRp sequence from accession MN851287.1 (from pooled organ tissues of wild bats) was not detected in these searches. Twenty-six other of the 115 sequences were found to derive from RefSeq accessions annotated as members of the genus *Victorivirus* and were included in the preceding analyses represented by Fig. 2. Thus, the 115 RdRp sequences included 72 not previously analyzed. MAFFT-L-INS-i, TrimAl, and IQ-TREE were then applied as before. In the resulting trees, the sequences from the 17 apparent eimerian viruses again consistently defined a discrete clade with strong branch support and with no other sequences identified as belonging to this clade (Fig. 3). These and the preceding results provide consistently strong evidence that the 18 apparent eimerian viruses, including the two new viruses reported here, define a distinct taxonomic unit, which we believe should be recognized as the genus “Eimeriavirus” in family *Totiviridae*, as also proposed by Xin et al. (2016). Though beyond the scope of this study and requiring further analysis, it appears from both Fig. 2 and Fig. 3 (a) that the genus *Victorivirus* may warrant subdivision into at least two new genera or subgenera, based on comprising at least two major subclades and (b) that other yet-to-be recognized genera may also be represented in these figures.

## DISCUSSION

The two newly identified viruses in this report are noteworthy in being derived from defined laboratory isolates of *Eimeria* species (*E. necatrix* isolate Yangzhou and *E. stiedai* isolate SCES/Sichuan), not from wild-collected samples that might contain mixtures of *Eimeria* strains or species. We therefore expect that the new sequences represent discrete virus strains of each respective species and isolate. Notably, both of these *Eimeria* host species are important causes of coccidiosis in food animals, *E. necatrix* in chickens and *E. stiedai* in rabbits.

Lee et al. have published a series of reports indicating the presence of a small RNA virus— genome size, ∼5.6 kbp; capsid diameter, ∼42 nm—in *E. necatrix* isolate Guelph (Lee et al., 1996; Lee and Fernando, 1998, 1999a, 1999b, 2000). Notably, in Lee et al. (1998), a partial sequence of 253 nt was reported from a cDNA clone of the purified genome of this virus. The sequence has not been deposited in GenBank but was copied from Lee et al. (1998) for use in this study. When compared with the sequence of EnecRV1-Yangzhou, using the local alignment program EMBOSS Matcher, this 253-nt sequence exhibited 78% identity and no gaps across its complete length. Moreover, when the 253-nt sequence was translated (no stop codons, 83 aa), it exhibited 83% identity and no gaps when compared with the RdRp sequence of EnecRV1-Yangzhou. These results suggest that the 5.6-kbp dsRNA identified by Lee et al. in *E. necatrix* isolate Guelph may represent another strain (EnecRV1-Guelph) of the same virus species as EnecRV1-Yangzhou, given their relatively high level of sequence identity.

As noted above, EstiRV1-SCES similarly appears to represent another strain of the same virus species as EstiRV1-TQCBD-12/0909, given their high level of sequence identity. Interestingly, EstiRV1-SCES lacks 20 nt at its 3′-terminus, but 117 nt at its 5′-terminus, relative to EstiRV1-TQCBD-12/0909 (NC_040530.1). To explain these missing residues, it is possible they are missing because reads from those terminal regions are simply absent from the SRA libraries from which the EstiRV1-SCES sequence was assembled. We found, however, that the extended 5′-terminal region of NC_040530.1 represents the reverse complement of a 112-nt region that spans residues 1357–1467 of NC_040530.1 (residues 1240–1350 in EstiRV1-SCES). Since such a long duplication is unusual, it seems possible that the 5′-terminal region unique to NC_040530.1 may reflect an error.

Both viruses newly reported in this study share the apparent stop/restart motif AUGA, where UGA would serve as the ORF1 (CP) stop codon and AUG would serve as the ORF2 (RdRp) start codon. An AUGA stop/restart motif between ORF1 and ORF2 is also found in EbruRV1 (NC_002701.1), EstiRV1-TQCBD-12/0909 (NC_040530.1), EtenRV1-JL610V (NC_026140.1), and Eimeria tenella RNA virus 1-like virus isolate EuB-TtV1 (MN851287.1). The partial sequence of Eimeria stiedai RNA virus 1 strain GUNCC_DN49375-6 (MT129770.1) does not cross the ORF1/ORF2 junction. Of the ten sequences representing putative eimerian viruses from feces of wild birds, six encompass the ORF1/ORF2 junction and have the following apparent stop/restart motifs: AUGA (MW826508.1), UGAUG (MZ218665.1), UAAUG (MZ218628.1, MZ218652.1, MZ218685.1), and UAAAUG (MW826406.1). Flexibility of relative positioning of the stop and start codons within the stop/restart motif, and even some allowance for greater separation of the two codons, is a feature of this mechanism as seen in victoriviruses and others (Firth and Brierley, 2012; Li et al., 2011). The only putative eimerian virus whose sequence crosses the ORF1/ORF2 junction but does not appear to have such overlapping or adjacent stop and start codons is Eimeria stiedai RNA virus 1 strain GUNCC_DN54384-11 (MT129769.1). This sequence, when translated, shows the UGA stop codon of ORF1 located 69 residues upstream of the putative AUG start codon of ORF2. In the GenBank accession itself, however, the authors have annotated ORF2 as though it starts with the CUG codon that overlaps the ORF1 stop codon, in the alternative stop/restart motif CUGA. This is an interesting possibility, which might be confirmed in the sequences of yet-to-be-discovered eimerian viruses. Use of a CUG start codon has been found in a number of other viruses and cellular organisms (Firth and Brierley, 2012; Touriol et al., 2003), including in the setting of a stop/restart motif (Lutterman and Meyers, 2007). A pseudoknot shortly upstream of the AUGA, etc., motif has been additionally implicated in the stop/restart mechanism of at least some victoriviruses (Li et al., 2011), but that element could not be readily identified in the eimerian viruses analyzed here.

Different virus species are commonly demarcated, albeit somewhat arbitrarily, by applying a threshold of maximum sequence identity; in other words, if two viruses have an identity score above that threshold, then they are commonly considered members of the same species. For members of genus *Victorivirus*, for example, that somewhat arbitrary threshold is currently set at 60% for the CP and RdRp proteins (Wickner et al., 2011). Having a coding-complete sequence for a particular virus is another criterion often applied before formally considering it to represent a new species. In this report, we have noted that there are eight apparent eimerian viruses for which coding-complete sequences have been reported to date, including the two new viruses reported here. Global pairwise comparisons of the CP and RdRp sequences of these eight viruses, obtained using EMBOSS Needleall, are shown in Fig. 4 and suggest that these eight viruses could be considered to represent seven distinct species. Only EstiRV1-TQCBD-12/0909 and EstiRV1-SCES would be clearly assigned as members of the same species, given their high level of sequence identity as also noted above. EbruRV1 and EnecRV1-Yangzhou approach a possible 60% threshold for species demarcation among the eimerian viruses, but because they derive from different *Eimeria* host species and are viruses for which horizontal transmission, especially between host species, is thought to be uncommon, we argue they should be considered to represent different virus species.

**Fig. 4.**
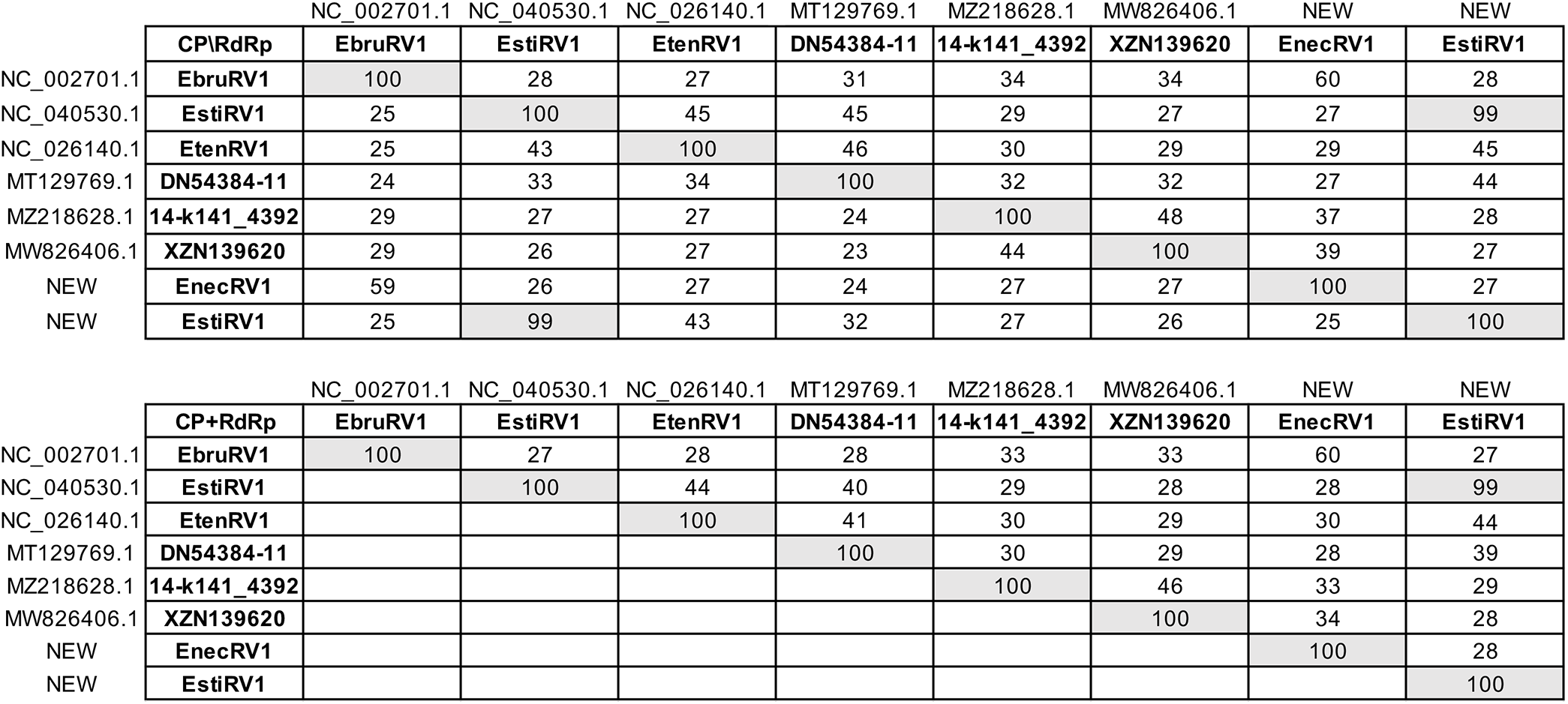
Pairwise scores. Identity scores (%) in pairwise global alignments were determined with EMBOSS Needleall using CP and RdRp sequences translated from the respective coding-complete nt sequences for putative eimerian viruses. RefSeq or GenBank accession numbers are provided for previously reported sequences. In the top grid, scores for CP are shown in the bottom left half, scores for RdRp in the top right half. In the bottom grid, scores for concatenated CP and RdRp sequences are shown. In addition to the high scores (100%) for self-comparisons, the high scores (99%) for comparisons between EstiRV1 NC_040530.1 (i.e., EstiRV1-TQCBD-12/0909) and EstiRV1 NEW (i.e., EstiRV1-SCES) are highlighted by shading.

As indicated in this report, viruses related to EnecRV1-Yangzhou and EstiRV1-SCES are found in a number of different *Eimeria* species, of which there are many in turn (∼1700 by one estimate; López-Osorio et al., 2020). In addition to evidence discussed above, for example, the literature includes other evidence for a related RNA virus in *E. nieschulzi* (Roditi et al., 1994; Sepp et al., 1991; genome size, ∼5.7 kbp; virus particle diameter, ∼39 nm). In fact, a 126-nt (42-aa) sequence reported from a cDNA clone of the purified genome of this virus is found in GenBank accession L25869.1 (Roditi et al., 1994), and a tBLASTn search showed its top hit (E-value, 1e-14; query cover, 100%; identity, 64%) to be MZ218671.1 (Totiviridae sp. isolate 13-k141_67720; Chen et al., 2022), which is one of the partial sequences of an apparent eimerian virus identified in Results (see Fig. 3). Indeed, the sequences of apparent eimerian viruses described above that were obtained from metagenomic samples (Chen et al., 2022; Kohl et al., 2021; Mahar et al., 2020; Zhu et al., 2022) might well represent viruses from yet other *Eimeria* species, as argued in Results. Based on Fig. 3, it seems possible to interpret that 10 or more species of eimerian viruses may be represented in that figure, perhaps derived from nearly as many *Eimeria* species. The prevalence of such viruses in members of the genus *Eimeria* begs the question of why that might be the case. It is interesting to speculate that these viruses, which are thought to mediate persistent, largely noncytopathic infections as do other *Totiviridae* members, play some beneficial role in eimerian biology that contributes to their prevalence, an idea that would seem useful to pursue in future work.

## ACKNOWLEDGMENTS

This work was supported by grants from the U.S. National Institutes of Health (No. R01 AI132445 to MLN), the National Natural Science Foundation of China (No. 32273028 to YX and No. 31972698 to JT), and the 111 Project D18007 and the Priority Academic Program Development (PAPD) of Jiangsu Higher Education Institutions (to JT).

